# Unusually slow spike frequency adaptation in deep cerebellar nuclei neurons preserves linear transformations on the sub-second time scale

**DOI:** 10.1101/2021.08.10.455865

**Authors:** Mehak M. Khan, Christopher H. Chen, Wade G. Regehr

**Affiliations:** Department of Neurobiology, Harvard Medical School, 220 Longwood Ave, Boston MA 02115

## Abstract

Purkinje cells (PCs) are spontaneously active neurons of the cerebellar cortex that inhibit glutamatergic projection neurons within the deep cerebellar nuclei (DCN) that in turn provide the primary cerebellar output. Brief reductions in PC firing rapidly increase DCN neuron firing. However, prolonged reductions in PC inhibition, as seen in some disease states, certain types of transgenic mice, and in acute slices of the cerebellum, do not evoke large sustained increases in DCN firing. Here we test whether there is a mechanism of spike-frequency adaptation in DCN neurons that could account for these properties. We find that prolonged optogenetic suppression of PC synapses *in vivo* transiently elevates PC firing that strongly adapts within ten seconds. We perform current-clamp recordings at near physiological temperature in acute brain slices to examine how DCN neurons respond to prolonged depolarizations. Adaptation in DCN neurons is exceptionally slow and bidirectional. A depolarizing current step evokes large initial increases in firing that decay to less than 20% of the initial increase within approximately ten seconds. Such slow adaptation could allow DCN neurons to adapt to prolonged changes in PC firing while maintaining their linear firing frequency-current relationship on subsecond time scales.

## Introduction

The cerebellum participates in motor tasks, motor learning, cognitive processing, social behaviors and many non-motor behaviors (Buckner, 2013; Reeber et al., 2013; Van Overwalle et al., 2014; Wang et al., 2014). To understand how the cerebellum contributes to these diverse functions, it is important to understand what controls cerebellar output. Purkinje cells (PCs) of the cerebellar cortex make strong GABAergic synapses onto glutamatergic projection neurons in the deep cerebllar nuclei (DCN) that in turn provide the output of the cerebellum. PCs provide an extraordinary amount of ongoing inhibition to the DCN. PCs fire spontaneously at high frequencies *in vivo* (Thach, 1968; Zhou et al., 2014a), and each DCN cell receives an estimated 400 Hz to 4 kHz of inhibition (40 PCs converging onto each DCN neuron firing at 10 to 100 Hz; Chan-Palay, 1973a, 1973b; Person and Raman, 2012). Yet, DCN cells are not silenced by this ongoing inhibition, and fire at basal rates of approximately 10-50 Hz *in vivo* and even higher rates during sensory or motor events (Eccles et al., 1974; LeDoux et al., 1998; McDevitt et al., 1987; Rowland and Jaeger, 2005; Rowland and Jaeger, 2008; Thach, 1975, 1970, 1968). Synchronous firing among PCs provides a means of transiently increasing and then suppressing inhibition to promote DCN firing (Person and Raman, 2012). Brief pauses in PC firing can also rapidly and transiently decrease inhibition to promote firing (Han et al., 2020).

Little is known about how DCN firing is regulated on long time scales, but there are reasons to suspect that DCN neurons slowly adapt to prolonged changes in PC firing. In PC degeneration (PCD) mutant mice where most PCs degenerate, it is expected that the loss of PC synapses would lead to disinhibition and elevated spiking in DCN neurons and in neurons of the vestibular nucleus (VN), another major output nucleus of the cerebellum (Bäurle et al. 1997). However, in PCD mice, the firing rates of VN neurons are not elevated, and rotation-evoked changes in VN neurons are much smaller than expected (Bäurle et al, 1997). This suggests that VN neurons may adapt their responses to the long-term absence of PC iputs. Similarly, in acute brain slices DCN neurons fire spontaneously at moderate frequencies, even though it is expected that the firing of DCN neurons would be substantially increased when inhibition from PCs is eliminated, because most PC axons in slice are severed (Jahnsen, 1986, Llinás and Mühlethaler, 1998, Aizenman et al., 1999, Czubayko et al., 2001, Uusisaari et al., 2007).

Together, these studies suggest that neurons targeted by PCs adapt to the prolonged absence of PCs. It is unknown if such an adaptive process is present, and if it is present, under what timescale it operates. This issue is also relevant for optogenetic manipulations of either PCs or DCN neurons to study behavior, where adaptation within the DCN might limit the ability of optogenetics to lead to sustained changes in DCN outputs. Adaptation could also be important during learning, because PCs undergo diverse forms of plasticity that can alter their excitability for hundreds of milliseconds to minutes (Belmeguenai et al., 2010; Gilbert, PFC, Thach, 1977; Grasselli et al., 2016; Heiney et al., 2014; Ito, Masao, Kano, 1982; Jirenhed et al., 2007; Rasmussen et al., 2008; Yang and Lisberger, 2014). It is therefore important to clarify the mechanisms of adaptation in DCN neurons.

Here we find that the sustained optogenetic suppression of PC synapses *in vivo* transiently elevates the firing of DCN neurons. The firing rate strongly adapts within approximately ten seconds, suggesting that there may be a mechanism of spike frequency adaptation in DCN neurons that acts on this timescale. We tested this possibility by examining the responses of DCN neurons to prolonged depolarizing current injections to determine if DCN neurons exhibit spike frequency adaptation. We found that a depolarizing current step evokes a large initial increase in firing that decays to less than 20% of the initial increase within approximately ten seconds. Such slow adaptation could allow DCN neurons to adapt to prolonged changes in PC firing, while maintaining DCN neurons’ linear firing frequency-current input relationship on subsecond time scales.

## Results

Previous studies in transgenic mice and brain slices suggested that the firing of DCN neurons could adapt within hours or days to the deprivation of PC inhibition. We used optogenetics to determine whether adaptation could also occur on a more rapid time scale. We used a transgenic approach to selectively express halorhodopsin in PCs (Pcp2-cre mice X eNpHR3.0-EYFP mice) (**Figure 1A**). We then positioned an optical fiber above the DCN to optically suppress PC to DCN axons and used an extracellular electrode to record responses of a DCN neuron. As expected, suppressing PC inhibition led to an increase in DCN spiking, but the effects of this supression were transient (**Figure 1B**). Initially, the average normalized firing frequency increased by 130%, but it eventually decayed to a 16% increase (**Figure 1C**). Although there was considerable cell-to-cell variability in the firing rate increases and the extent of adaptation (**Figure 1DE**), it was clear that the ability of this optogenetic approach to increase the firing of DC neurons in a sustained manner was limited. This adaptation of normalized firing was approximated by an exponential decay with a time constant of 8.3 s. It is unclear whether such an adaptation in firing rate refects limitations in optical suppression of PC axons or the adaptation of DCN neuron firing to a sustained reduction of PC inhibition.

**Figure 1.**
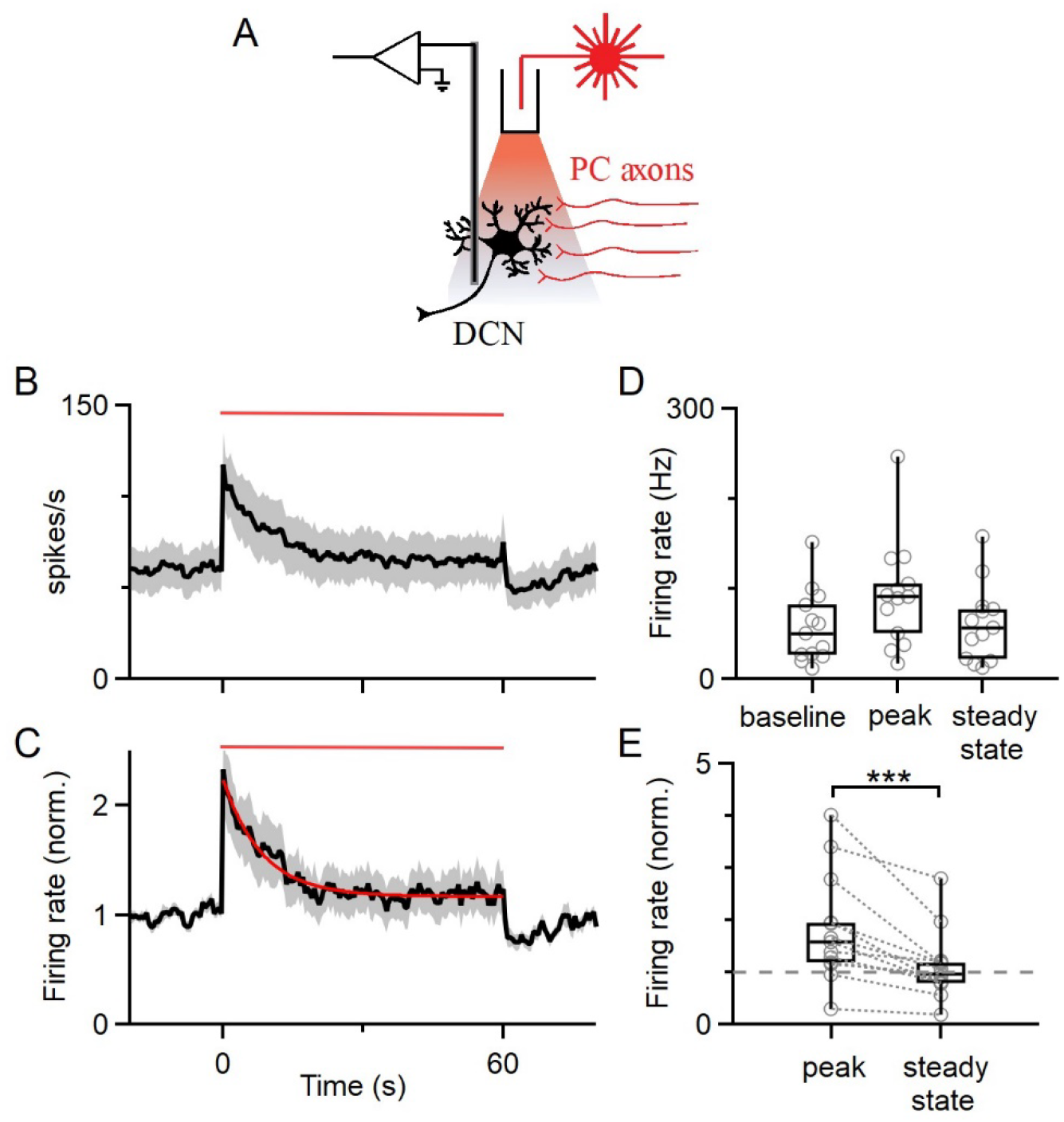
DCN neurons slowly adapt to sustained decreases in Purkinje cell inhibition in vivo. A. Schematic showing recording configuration where the axons of halorhodopsin-expressing PCs were optogenetically inhibited with light through an optical fiber placed in the DCN. B. The average effects of 60 s optical inhibition of PC axons (red bar) on the average firing of DCN neurons. Standard error is shaded in gray. C. The normalized firing of DCN neurons in B. Standard error is shaded in gray. D. The average firing rate of each DCN neuron is plotted for baseline firing, during the initial period of PC axon inhibition, and for the steady state firing reached near the end of steady state inhibition. E. Normalized firing rates for the data in D. Normalized peak and steady-state firing rates are joined by a dotted line for each neuron.

Although previous studies have examined spike frequency adaptation of DCN neurons within hundreds of milliseconds to several seconds (Jahnsen, 1986, Llinás and Mühlethaler, 1998, Aizenman et al., 1999, Czubayko et al., 2001, Uusisaari et al., 2007), the properties of spike frequency adaptation on the tens of seconds time scales are not known. We therefore tested the possibility that DCN neurons adapt their firing in a manner that could contribute to the adaptation of firing observed in **Figure 1**. Injecting a depolarizing current is a practical way to examine spike frequency adaptation in brain slice. To study spike frequency adaptation of DCN neurons, we recorded in whole-cell current clamp from DCN neurons at near physiological temperatures (36°C) in acute slices from juvenile (P25 to P40) mice and injected constant depolarizing currents for 50 s. We recorded from large DCN neurons that correspond to glutamatergic projection neurons (Uusisaari et al., 2007; Turecek et al. 2016). DCN neurons initially fired at high frequencies, and their rate of firing adapted very slowly (**Figure 2**). This is shown for an example cell that initially fired at 180 Hz and gradually slowed to ~ 23 spikes/s (**Figure 2A**). We defined the ratio of steady-state to peak firing as the adaptation ratio (steady-state firing rate/peak firing rate, ss/peak) and the decay time (t_decay_) as the time-constant of an exponential function best fit to the firing rate response. In **Figure 2A** the adaptation ratio is 0.13 and the decay time is 7.4 s. There was variability in the time-course and extent of adaptation (**Figure 2B, D**), with an average adaptation ratio of 0.17 ± 0.02 and an average decay time of 4.9 ± 0.4 s (**Figure 2C**).

**Figure 2:**
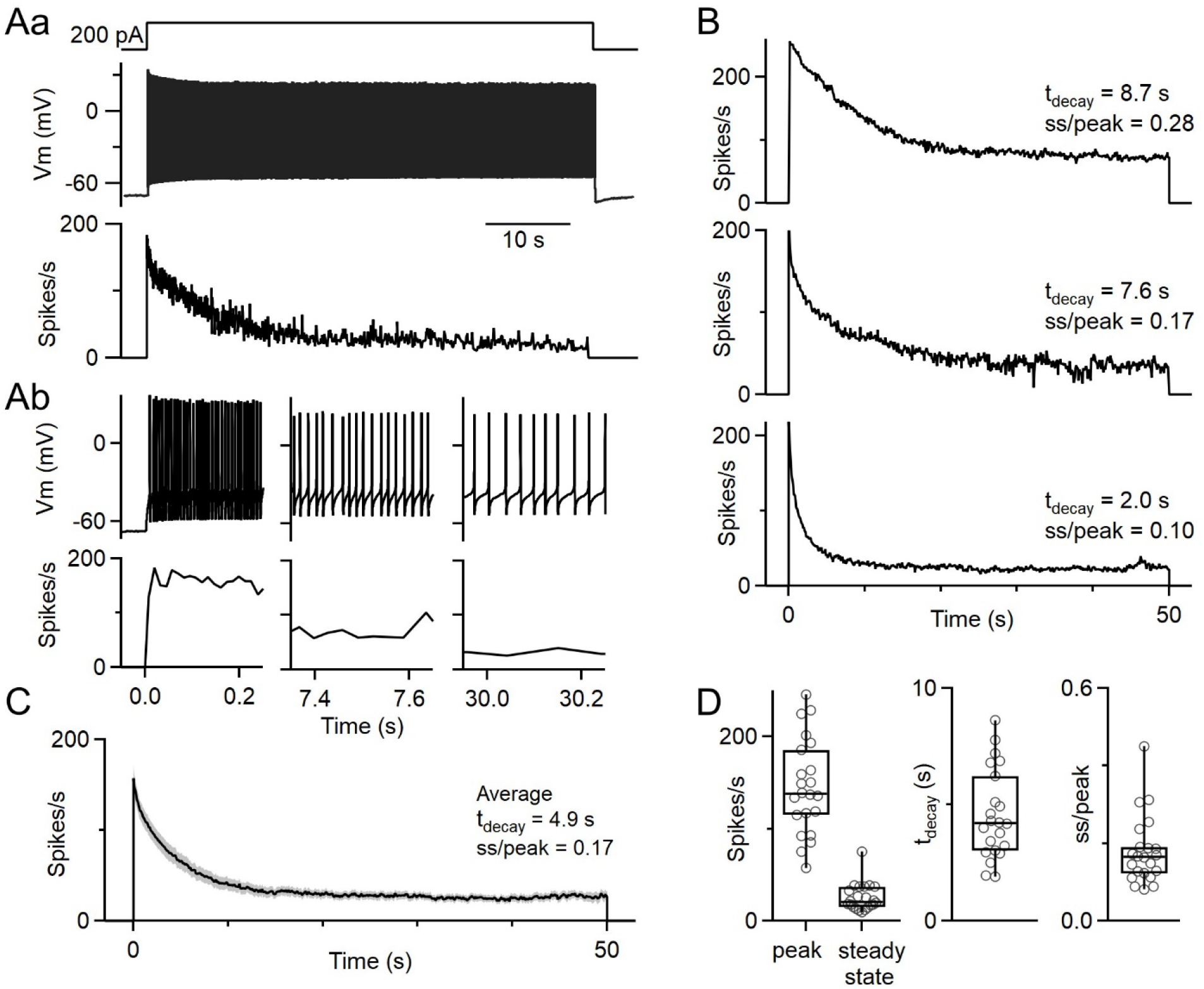
DCN neurons exhibit robust, slow spike frequency adaptation. A. a. Representative cell response to a 50 s depolarizing current step (200 pA) delivered to a DCN cell in whole-cell current-clamp (top), the resulting voltage trace (middle), and the instantaneous firing frequency of spiking response (bottom). b. Expanded views of different periods of the voltage trace in Aa are shown (top), and the corresponding instantaneous firing frequencies (bottom). Expanded views of the same time points as immediately above for instantaneous firing rate. B. Example instantaneous firing frequencies for three different DCN cells that were depolarized for 50 s. Decay times and adaptation were determined as above. Decay times were determined from an exponential fit to the instantaneous firing frequency during the depolarization. Adaptation ratio (ss/peak) was calculated as the steady-state (ss) firing rate divided by the peak firing rate (peak). C. Average response to a 50 s current step for n = 22 DCN neurons (7 animals), with the standard error overlaid and shaded in gray. D. Summary data for the peak and steady-state firing rates, decay times, and adaptation ratios (N=22).

In many neurons, spike-frequency adaptation is thought to reflect the build up of a hyperpolarizing conductance that suppresses spiking and also leads to an afterhyperpolarization (AHP) after spiking is terminated (Schwindt et al., 1988, Foehring et al, 1989, Pedarzani and Storm, 1993, La Camera, 2006, Nishimura et al, 1989, Sawczuk et al, 1995, Safronov and Vogel, 1996, Madison and Nicoll, 1982, 1984, 1986, Ha et al, 2016, Faber and Sah, 2002, Khawaja et al, 2007, Barazza et al., 2009, Sanchez-Vives et al., 2000a). We found that a prominent AHP was observed in DCN neurons after the cessation of spiking (**Figure 3A**). To examine the relationship between the magnitude of the AHP and the extent of adaptation, we delivered depolarizing current steps with a range of durations (0.3 s to 30 s).

**Figure 3:**
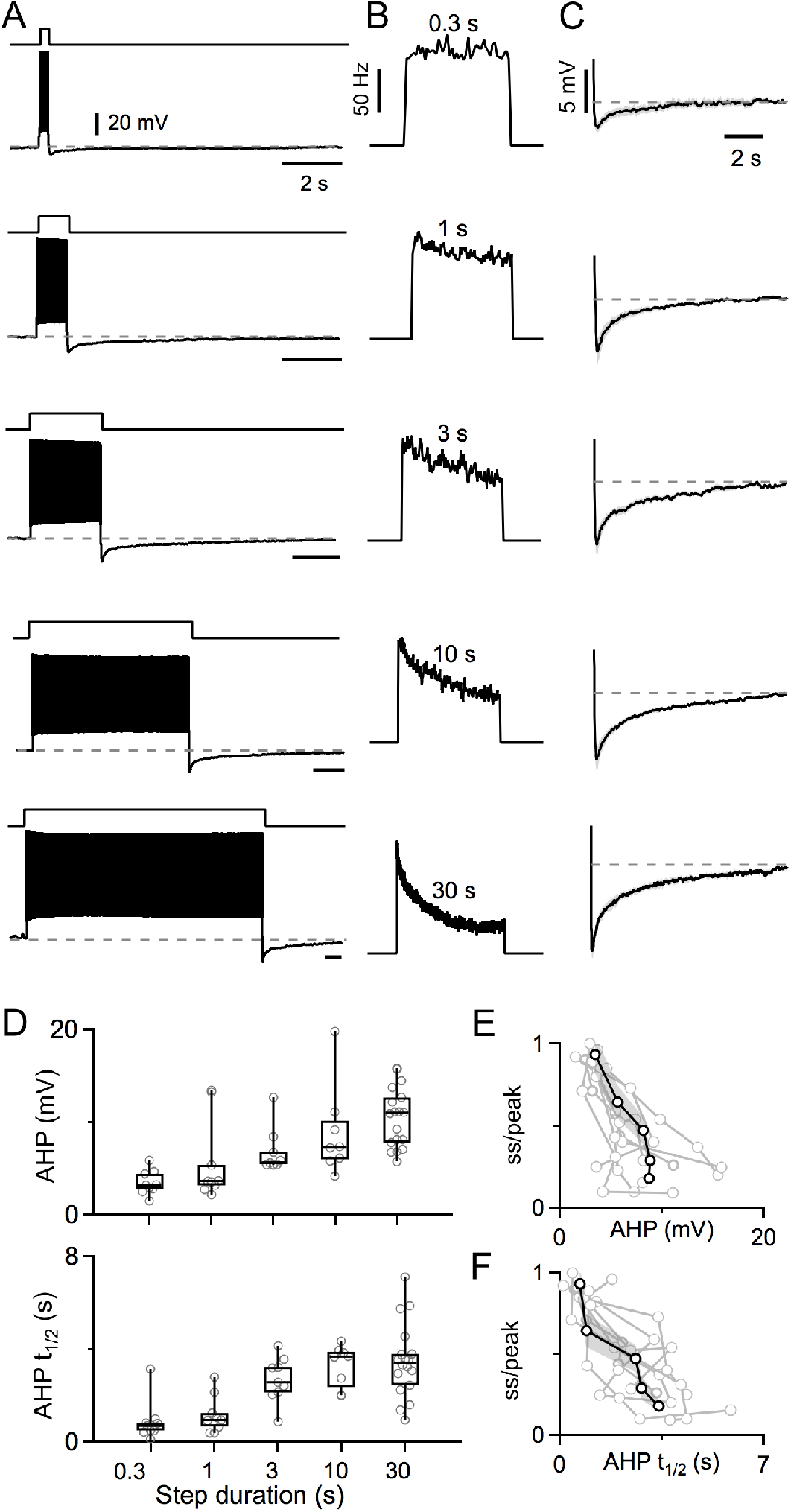
Prolonged spiking leads to a slow afterhyperpolarization that is dependent on spiking duration. A. Firing and afterhyperpolarization for a DCN neuron are shown in response to delivering depolarizing current steps of varying duration. B. Instantaneous firing frequencies are shown for the spiking shown in A. C. Voltage traces showing the average afterhyperpolarizations (AHPs) following 0.3 s, 1 s, 3 s, 10 s, and 30 s current steps (N = 7 cells from 5 animals, standard errors in gray). D. Summaries of the amplitudes and half-recovery times of the AHPs for individual cells (circles) after current steps of the indicated durations. Half-recovery time was calculated as half the time for the AHP to return to the initial membrane potential before the step. N = 7 cells (5 animals). E. Summary of firing rate at the end of a depolarizing current step lasting 0.3 s to 30 s divided by the peak firing of the step (ss/peak), as a function of the magnitude of the subsequent AHP. Responses from individual neurons are shown (light grey), with the average response overlaid in black and standard error shaded in dark grey. F. Same as in E, but ss/peak is plotted as a function of the AHP half-recovery time.

During 0.3 s steps, the extent of adaptation and the amplitude of the AHP were both small (**Figure 3A-B, top**). As the durations of current steps were increased, adaptation became more prominent (**Figure 3B**) and the amplitude of the AHP increased (**Figure 3C-D**). Longer current steps were also followed by longer-lasting AHPs, as measured by the time taken for the AHP to return halfway to the initial resting potential, which is the AHP half-recovery time (AHP t_1/2_) (**Figure 3CDF**). We observed a strong negative correlation between the amplitude of the AHP and the firing rate at the end of the depolarizing step (Pearson’s correlation coefficient, ρ = − 0.641), and for AHP t_1/2_ versus firing rate at the end of the depolarizing step (ρ = −0.701).

Spike frequency adaptation and afterhyperpolarizations have been shown to be mediated by a variety of mechanisms in other types of neurons, and we tested several possibilities for DCN neurons (**Figure 3-figure supplement 1, Figure 3-figure supplement 2**). One possibility was that high frequency firing leads to the buildup of intracellular calcium and the activation of calcium-dependent conductances. We found, however, that spike frequency adaptation remained prominent in low external calcium and in 0 external calcium with 1 mM EGTA added to reduce extracellular calcium to very low levels. We also found that spike frequency adaptation was unaltered by apamin, a blocker of calcium-activated SK channels, or by XE991, a blocker of M-current (Kv7, KCNQ channels). Finally, we found that spike frequency adaptation was intact in SLICK/SLACK double knockout animals in which sodium-activated potassium channels are eliminated. The amplitude of the AHP was reduced in low external calcium, attenuated when external calcium was eliminated, and unaffected by a either an SK or by an M-current blocker, or in SLICK/SLACK double KO mice. These findings suggest that calcium-activated channels, SK channels, KCNQ channels, and sodium-activated potassium channels alone do not mediate spike-frequency adaptation or the AHP. Further studies are needed to clarify the mechanisms of spike-frequency adaptaion in DCN neurons.

The slow AHP we observe has the potential to reduce excitability following high-frequency spiking and could also contribute to spike frequency adaptation. We examined this possibility by using a current injection protocol consisting of a 200 pA current injection, followed by a 400 pA current step, and then returning to a 200 pA current (**Figure 4**). As shown for an example cell, the initial depolarization transiently elevated spiking that then decayed to a steady-state level, and a further increase in depolarization elevated spiking that again adapted to a new steady-state level (**Figure 4A**). In response to the additional depolarization, steady-state spiking was elevated (11.5 spikes/s for 400 pA vs. 5.4 spikes/s for 200 pA), and the time-constant of spike adaptation was slower (6.3 s for t_decay2_ vs. 3.8 s for t_decay1_). Although the reason for the differences in the time courses of adaptation are unclear, this observation indicates that in addition to differences in adaptation between cells, an individual cell can exhibit different time courses of adaptation. After stepping down from 400 pA to 200 pA, spiking stopped for 13.8 s (t_firing_) (**Figure 4A**). Seven cells and their average responses showed similar trends (**Figure 4B-D**). This result indicates that adaptation in DCN neurons is bidrectional. DCN neuron firing slowly adapts during periods of elevated firing and recovers from reduced firing in a manner that is consistent with the time-course of the long-lasting conductance that mediates the AHP.

**Figure 4:**
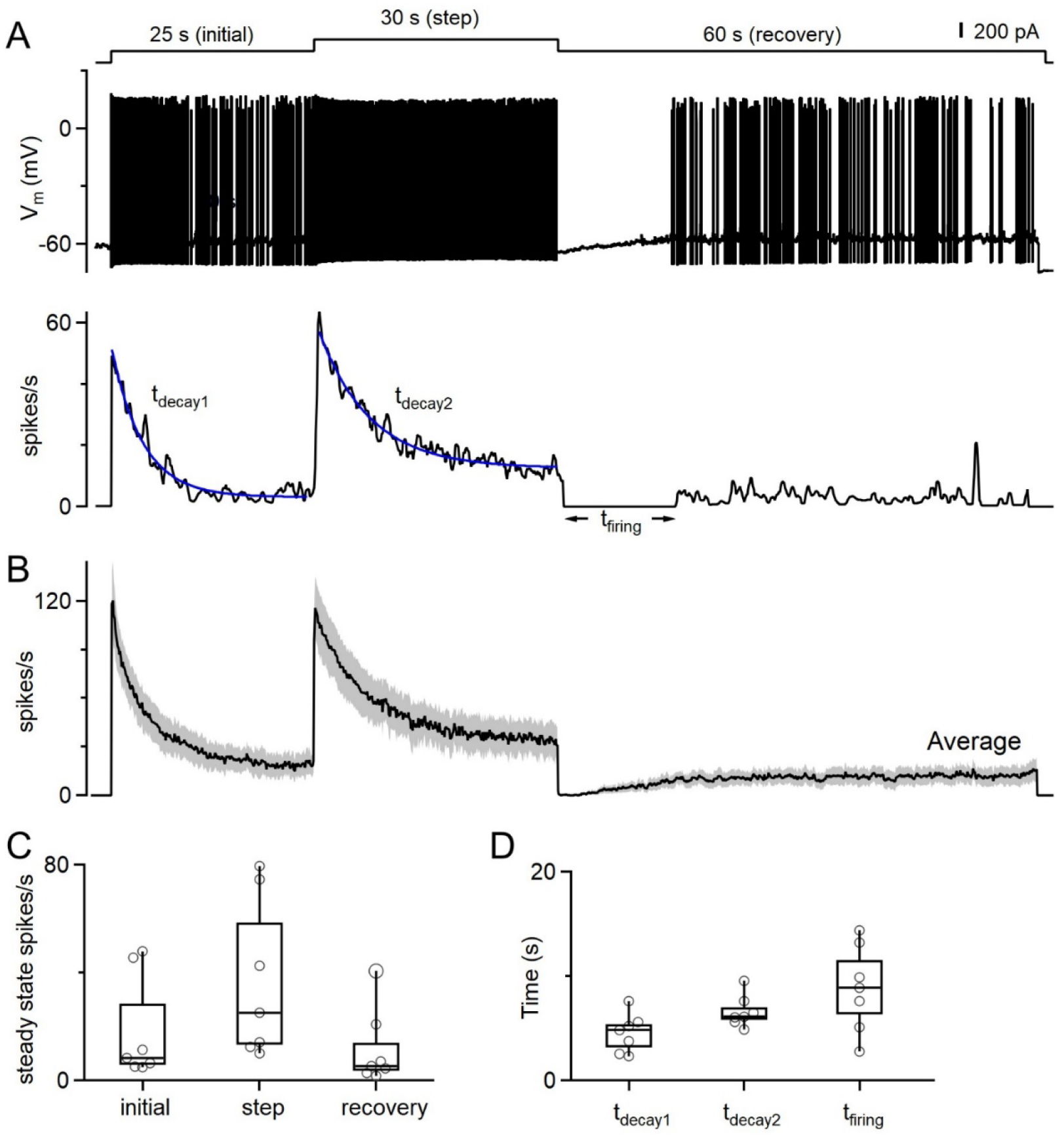
DCN firing bidirectionally adapts to increases and decreases in depolarization over seconds. A. Top: A three-step depolarizing current injection protocol delivered to a DCN cell, starting with an initial 200 pA depolarization for 25 s, followed by a step to 400 pA for 30 s, and then a 200 pA depolarization for 60 s. Middle: Voltage response during a current-clamp recording from a DCN neuron Bottom: Instantaneous firing response. B. Average instantaneous firing frequency evoked by the three-step current injection protocol. N = 7 cells (5 animals). C. Summary data for the steady-state firing at the ends of the initial depolarization, the 400 pA step, and for the 60 s recovery. D. Summary data for the decay-times to reach steady state for the initial current injection (t_decay1_), and for the 400 pA step (t_decay2_). Decay times were determined from an exponential fit to the instantaneous firing frequency during the indicated period. In all cases the spiking stopped when the depolarization was decreased from 400 pA to 200 pA, and the time taken for firing to resume is summarized for each cell (t_firing_).

The slow adaptation of DCN neurons and their lack of adaptation on rapid time scales suggests that DCN neurons respond differentially to brief and long-lasting depolarizations. We examined this possibility by delivering a series of depolarizing current steps of varying duration to DCN neurons. Responses are shown for sequential 1 s depolarizations of 50, 100, 150, 200, 150, 100, and 50 pA (**Figure 5A**, *top*). Spiking is shown for the initial 100 ms (*black*) and the final 100 ms (*blue*) of each depolarization. DCN firing rates varied with the magnitude of the current steps (**Figure 5A** *top***, B** *left*). Firing rates were similar for the first and last 100 ms (**Figure 5A**, top), consistent with minimal adaptation. Responses were very different when the duration of each step was 30 s (**Figure 5A**, *bottom*). For step increases in the depolarizing current, the initial firing was much higher at the start of the depolarization than at the end, but when depolarizing currents were decreased, the initial firing was much lower at the start than at the end (**Figure 5A** *bottom*, **B** *right*). Similar experiments were performed for 0.3 s, 1s, 3 s, 10 s and 30 s steps for a number of DCN neurons, with average responses as a function of time (**Figure 5C**, left) and the average firing rate in the initial 100 ms (black) and the final 100 ms (blue) plotted for each current step (**Figure 5C**, *right*). For 0.3 s to 1 s steps, the firing of DCN neurons tracked the magnitude of the depolarization (**Figure 5C**, *top*). Responses to steps that were 1 s long most faithfully tracked the change in depolarization over time. For 3 s steps, a small amount of adaptation is apparent and there is a small decrease in the firing rate by the end of the current step. Longer steps resulted in more adaptation and increasing divergence between initial and steady-state firing (**Figure 5C**). Overall, these results indicate that for short depolarizing current steps, DCN neurons faithfully convert the magnitude of the depolarizing current into a spike rate. For long changes in the magnitude of depolarizing inputs, DCN neurons slowly adapt and remain responsive to additional changes in the magnitude of depolarizing currents.

**Figure 5:**
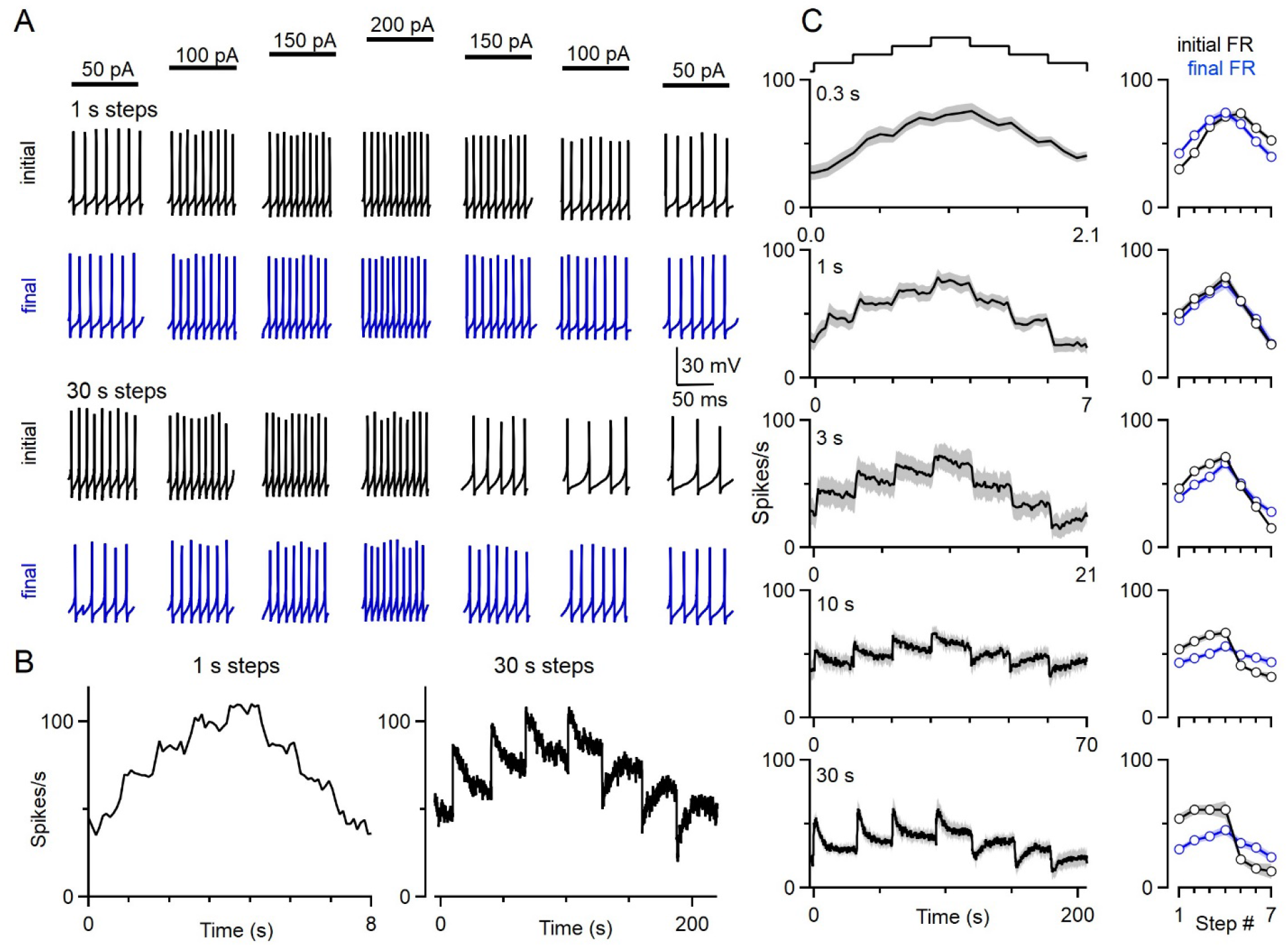
The firing of DCN neurons evoked by a series of current steps. A. Response of a representative DCN cell to sequential depolarizing current steps. The initial 100 ms (black) and the final 100 ms (blue) are shown for each current step. Before delivering the sequential current steps, DCN cells were firing at a steady-state level in response to a 100 pA current injected beforehand (at t = −20 s relative to start of the sequential current steps) to maintain baseline firing. B. Instantaneous firing frequencies evoked by 1 s (left) and 30 s (right) current steps are shown for the cell in A. C. Summary of the firing responses to current steps of varying duration. Plots on the left show the average instantaneous firing frequency of DCN cells in response to the depolarizing current ramp where each step is of the indicated duration. Standard error is shaded in grey. Plots on the right show the average firing rates for the initial (black) and final (blue) 100 ms (N = 7 cells, N = 6 animals).

## Discussion

Our main finding is that DCN neurons exhibit a robust, remarkably slow form of spike-frequency adaptation. This adaptation is so slow that it does not impact responses on the hundreds of milliseconds or several seconds time scales, and DCN neurons are able to respond linearly to changes in injected current on these short time scales. However, on longer time scales, DCN neurons adapt their firing in a manner that allows them to remain responsive to subsequent changes in excitatory or inhibitory drive.

### Previous studies of DCN neurons

There have been extensive studies of DCN neurons, but there have been no previous reports of the slow spike frequency adaptation we report here. This is likely because of differences in experimental conditions and the experimental approach. Many previous studies used short depolarizations of hundreds of milliseconds (Jahnsen, 1986, Llinás and Mühlethaler, 1998, Aizenman et al., 1999, Czubayko et al., 2001, Uusisaari et al., 2007, Witter and De Zeeuw 2016, Han and Turner 2014), and the slow spike frequency adaption we see is not apparent during such brief steps (**Figure 2Ab**, left). Numerous studies also focused on much younger animals than those studied here (Aizenman et al., 1999, Raman et al. 2000, Czubayko et al., 2001, Uusisaari et al. 2007, Zhang and Raman 2009), and adaptation may have different properties in young animals. Careful examination of DCN spiking evoked by somewhat more prolonged depolarizations in other studies are consistent with the slow adaptation we observe. In young rats (P16), spike frequency adaption of about 20% occurred for a 5 s depolarization (Czubayko et al., 2001). In more mature guinnea pigs, a 2 s depolarization resulted in highly variable adaptation that ranged from a 50% reduction in firing rate to no reduction at all (Jahnsen, 1986). Although the time course of adaptation was not characterized in that study, it is likely that a 2 s step was insufficiently long to reveal the full extent of spike frequency adaptation. Thus, previous studies did not use protocols that were suited to studying the slow spike-frequency adaptation pesent in DCN neurons.

### Implications for the cerebellar processing

The properties of spike-frequency adaptation of DCN neurons has several implications for cerebellar processing. We found that on average DCN firing adapts to 17± 2% of initial firing with a decay time of 4.9 ± 0.40 s (**Figure 2**), that DCN adaptation is also bidirectional (**Figure 4, Figure 5**), and that adapted cells remain remarkably sensitive to further changes in depolarization (**Figure 4, Figure 5**). On short time-scales that are less than 1 s, DCN firing reliably encodes changes in PC firing. This would ensure that cerebellar output could preserve signals generated from short-term changes in PC activity, such as those occurring during fine movements or error signals generated on a sub-second timescale. However, adaptation makes the DCN less sensitive to slower, more prolonged changes in input activity. PCs are known to undergo long periods of modulation in their excitability and activity, such as during motor learning (Belmeguenai et al., 2010; Gilbert et al., 1977; Grasselli et al., 2016; Heiney et al., 2014; Ito, Masao, Kano, 1982; Jirenhed et al., 2007; Rasmussen et al., 2008; Yang and Lisberger, 2014). In these situations, DCN cells could initially generate large responses but gradually adapt to maintain a stable firing range. Without such adaptation, DCN spiking could saturate and these neurons might be unable to respond to further changes in PC activity. When PC firing rates are increased, the adaptation of firing rates described here will act in concert with the slow depression of the PC to DCN synapse (Pedroarena 2020).

Spike-frequency adaptation of DCN neurons is well-suited to protecting against the loss of PC inhibition. Many types of ataxia involve a long-term decrease in PC inhibition of the DCN, often as a result of chronic decreases in PC firing, decreases in the strength of PC synapses, or PC death. PCs are also susceptible to hypoxia (Au et al., 2015; Welsh et al., 2002). Without spike frequency adaptation, a strong reduction of PC inhibition is expected to lead to extremely high sustained firing of DCN neurons that would likely result in the death of DCN neurons, thereby compromising the ability of the cerebellum to influence the rest of the brain. However, DCN neurons can survive in these mouse models despite strong reductions in PC inhibition, and spike-frequency adaptation likely helps to prevent large sustained increases in DCN firing. It is clear that spike-frequency adaptation cannot overcome profound deficits in PC inhibition and spare all cerebellar-dependent behaviors, as is apparent in motor deficits that accompany PC loss (Catterall et al., 2010, 2008; Cook et al., 2020; Levin et al., 2006; Tsai et al., 2018; Walter et al., 2006, Bosch et al., 2015; Dougherty et al., 2013, 2012; Hourez et al., 2011; Hurlock et al., 2008; Jayabal et al., 2016; Kordasiewicz and Gomez, 2007; Larivière et al., 2019; Shakkottai et al., 2011, 2009; Tsai et al., 2018). It is also likely that in brain slice, spike frequency adaptation prevents DCN neurons from firing at high rates in the absence of PC inputs. Finally, spike frequency adaptation likely contributes to the observation that acutely increasing PC activity or decreasing DCN activity reduces aggression (Heath 1977, Cooper et al. 1976, Reis et al. 1973, Jackman et al. 2020, Zanchelli and Zoccolini, 1954), whereas chronically removing PC inhibition by lesioning the cerebellar cortex, which would be predicted to increase DCN firing, similarly reduces aggression (Sprague and Chambers 1959, Berman 1997).

### Implications for the manipulation of DCN activity

The bidirectional spike frequency adaptation we observe will oppose conventional attempts at manipulating the activity of DCN neurons. It is likely that spike-frequency adaptation in the DCN conributes to the transient effects of sustained optogenetic suppression of PC axons on the firing of DCN neurons (**Figure 1**). Spike frequency adaptation could also limit the efficacy of sustained deep brain stimulation of the cerebellar nuclei, which is a promising approach for treating dystonia, ataxia, essential tremor and a variety of other neurological and neuropsychiatric disorders (Miterko et al., 2019).

### Comparison to spike frequency adaptation AHP in other cell types

Spike-frequency adaptation and AHPs have been described in many types of neurons. In most cases the time scales are much faster than we observed in DCN neurons, lasting tens or hundreds of milliseconds (Schwindt et al., 1988, 1989, Foehring et al, 1989, Pedarzani and Storm, 1993, Fleidervish et al, 1996, Pineda et al, 1999, Higle and Contreras, 2006, Nishimura et al, 1989, Sawczuk et al, 1995, Safronov and Vogel, 1996, Madison and Nicoll, 1982, 1984, 1986, Tzingounis et al, 2007, Ha et al, 2016, Sacco et al, 2003, Bischoff et al, 1998, Faber and Sah, 2002, Khawaja et al, 2007, Hardman and Forsythe, 2009). There are several types of cells that exhibit slow spike frequency adaptation: visual cortical neurons, subthalamic nucleus neurons, and fast-spiking GABAergic cortical interneurons (Barraza et al., 2009; Descalzo et al., 2005; La Camera et al., 2006; Sanchez-Vives et al., 2000a, 2000b). In V1 neurons, slow spike-frequency adapation allows neurons to adapt to constant stimuli, while preserving their ability to respond to rapidly changing stimuli (Sanchez-Vives et al. 2000ab). In subthalamic nuclei, it has been proposed that slow adaptation removes slow trends in the average rates of inputs (Barraza et al. 2009). Major questions remain regarding the mechanisms of slow spike frequency adaptation in all types of neurons where this phenomenon has been observed. The mechanism responsible for slow adaptation of FS neurons is not known (La Camera et al, 2006). In V1 neurons, an AHP produced by a K^+^ current is implicated, and it has been proposed that a build up Ca^2+^ and/or Na^+^ may activate this current (Sanchez-Vives et al. 2000ab), and Na^+^-activated K^+^ channels were the favored mediators. In subthalamic nuclei neurons, a K^+^ current is also implicated, but it is thought to be independent of Ca^2+^ and Na^+^ (Barraza et al. 2009). For DCN neurons we also think that a hyperpolarizing current is involved, but we were unable to eliminate adaptation by eliminating Ca^2+^ entry, by eliminating Na^+^-activated K^+^ channels in SLICK/SLACK KO mice, by blocking M current, or by blocking SK Ca^2+^-activated K^+^ channels. It remains possible that these different types of neurons share a common mechanism of spike frequency adaptation that has yet to be determined. It is also likely that slow spike adaptation is more common than is currently appreciated. Just as we revealed slow adaptation here in DCN neurons by using prolonged depolarizations, it is likely that extending the duration of depolarization in other cell types will reveal slow adaptation in some other types of cells.

## Methods

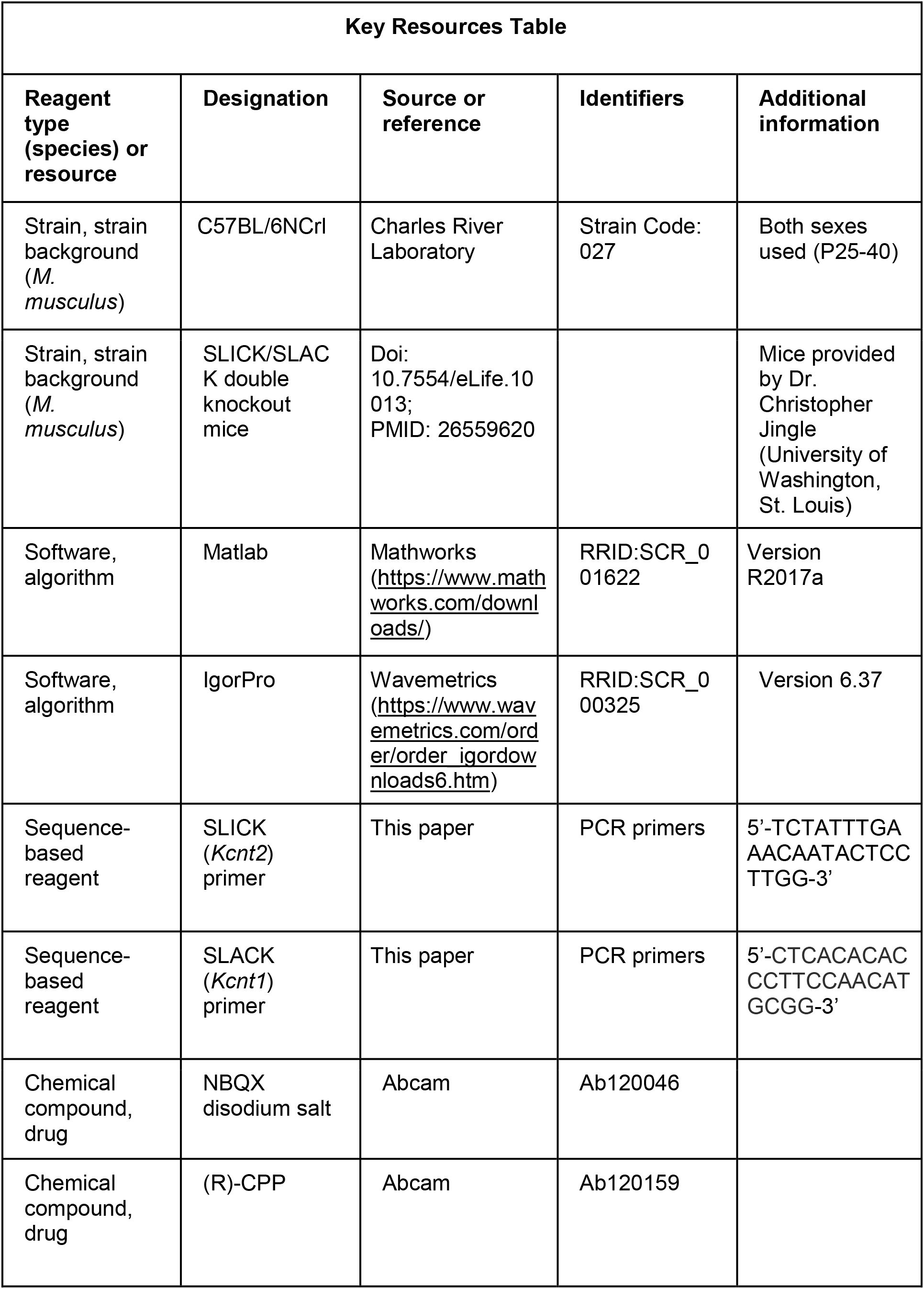
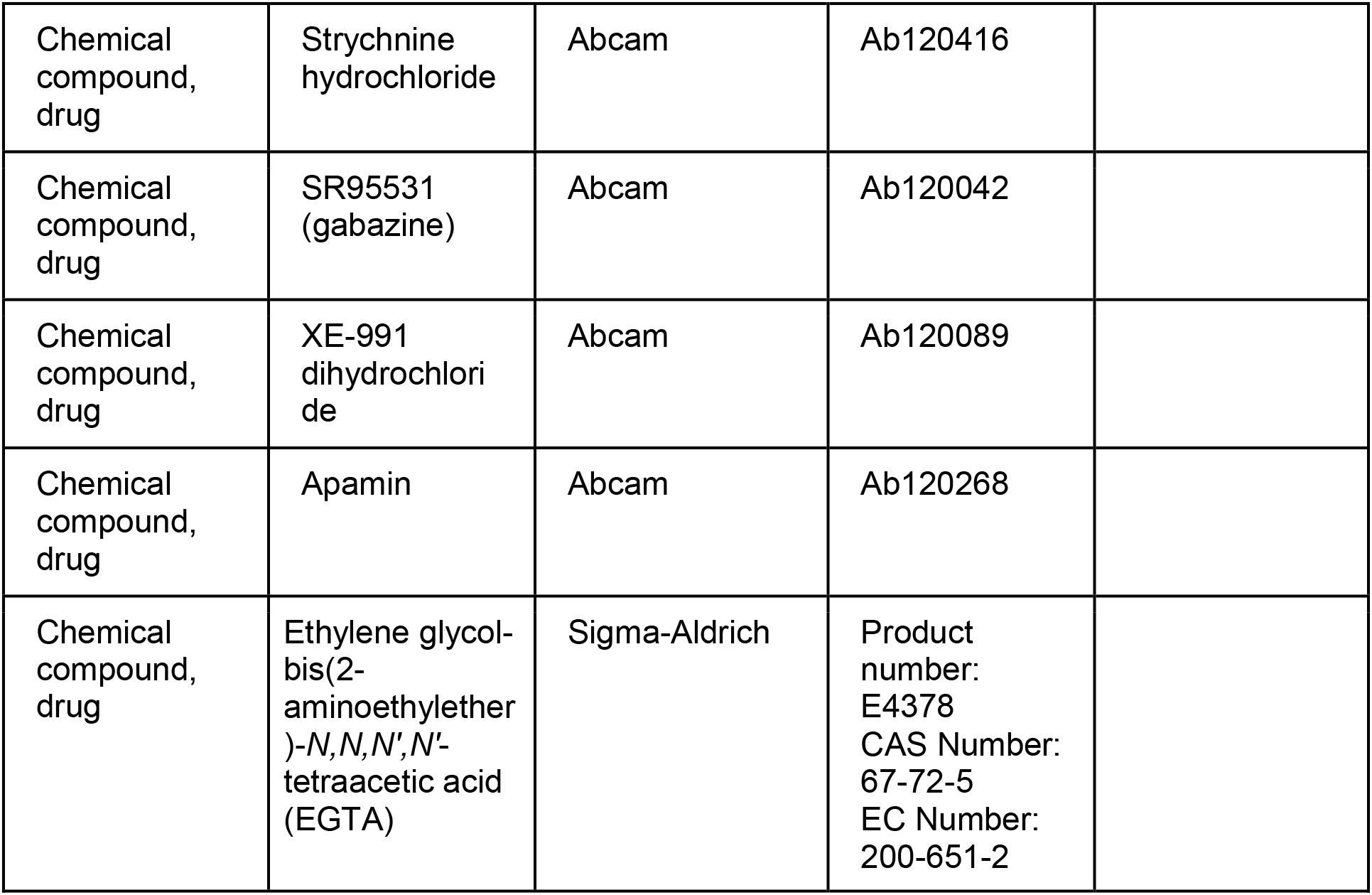

### Ethics

All animal procedures were carried out in accordance with the NIH and Animal Care and Use Committee (IACUC) guidelines and protocols approved by the Harvard Medical Area Standing Committee on Animals (animal protocol #1493).

### Animals

P25-40 C57BL/6 wildtype mice (Charles River Laboratories) of both sexes were used for most acute slice experiments. In **Figure 3 – figure supplement 1**, P25-40 SLICK/SLACK double knockout (dKO) mice of both sexes were used where indicated. For in vivo experiments, we used Pcp2-Cre mice (Jackson Laboratory, 010536) crossed with eNpHR3.0EYFP (Halo) mice (Ai39; Jackson Laboratory, 014539) to optogenetically inhibit PC firing.

### In Vivo Electrophysiology

*In vivo* recordings from PCs and DCN neurons were made from awake, head-restrained Pcp2-cre mice crossed to eNpHR3.0-EYFP (Halo) mice using an acutely inserted silicon probe (Cambridge Neurotech, Cambridge, England). To prepare for recordings, mice were anesthetized with 2% isoflurane and implanted with a head-restraint bracket using metabond (Parkell, Edgewood, NY). A craniotomy (~1 mm diameter) was made over the vermis (1-2 mm on either side of the midline and - 6.8mm posterior to bregma to target the fastigial and parts of the interposed nuclei). Craniotomies were sealed with kwik-sil (World Precision Instruments, Sarasota, FL) until the day of the recording. After surgery, mice were given the analgesic buprenorphine. All animals were acclimated to head-restraint on a freely-rotating cylindrical treadmill for at least one session prior to the recording session.

For experiments involving inhibition of PC axons, mice underwent the initial procedure above. The silicon probe used for recording included an attached optical fiber (200 μm diameter, 0.39 NA, Thorlabs or with Lambda-B fiber attach (100 μm diameter, 0.9 mm taper; Cambridge Neurotech)). Light was delivered directly through the attached optical fiber, in pulses as indicated in the text. PCs were identified by the presence of complex spikes and responsiveness to light. DCN were identified by their depth, response to light, and absence of complex spikes.

### Analysis of In vivo electrophysiology

Data were acquired at 20 kS/s (1 Hz to 10 kHz bandpass) using an Intan acquisition system (Intantech). Spikes were sorted in Offline Sorter (Plexon), and analyzed in Matlab (Mathworks, MA).

### Slice preparation

Mice were anesthetized with ketamine / xylazine / acepromazine and transcardially perfused with warm (34° Cel) choline-ACSF solution containing (mM): 110 Choline Cl, 2.5 KCl, 1.25 NaH_2_PO_4_, 25 NaHCO_3_, 25 glucose, 0.5 CaCl_2_, 7 MgCl_2_, 3.1 Na-Pyruvate, 11.6 Na-Ascorbate, and 5 μ M NBQX and 2.5 μM (R)-CPP, oxygenated with 95% O2 / 5% CO2. To prepare sagittal cerebellar slices, the hindbrain was removed, a cut was made down the midline of the cerebellum, and the halves of the cerebellum were glued down to the slicing chamber. 170 μm thick sagittal slices were cut with a Leica 1200S vibratome in warm choline-ACSF. Slices were transferred to ACSF solution containing, in mM: 127 NaCl, 2.5 KCl, 1.25 NaH_2_PO_4_, 25 NaHCO_3_, 25 glucose, 1.5 CaCl_2_, and 1 MgCl_2_ maintained at 34°C for 10-12 minutes before being moved to room temperature for 20-30 minutes before beginning recordings.

### Electrophysiology

Whole-cell voltage current-clamp recordings were performed on spontaneously firing, large diameter (20-25 μm) neurons in the lateral and interposed deep cerebellar nuclei. Borosilicate glass electrodes were filled with an internal solution containing, in mM, 150 K-Gluconate, 3 KCl, 10 HEPES, 0.5 EGTA, 3 Mg-ATP, 0.5 Na-GTP, 5 Phosphocreatine-tris2, and 5 Phosphocreatine-Na_2_ adjusted to pH 7.2 with KOH. The osmolarity was adjusted to 300 mOsm. High resistance (3-4 MΩ) electrodes were used to minimize dilution of cytosolic components, and only recordings where series resistance was below 15 MΩ were accepted. Series resistance was compensated up to 80% for the estimated capacitance of the cell body (5 pF). In most all cases, DCN neurons spontaneously fired upon breaking in at voltages between −50 to −60 mV. Hyperpolarizing current was injected to keep cells from spontaneously spiking between experimental trials. As a result, cells typically “rested” between −65 to −75 mV. Reported voltages were corrected for a liquid junction potential of 10 mV between the internal solution and ACSF bath solution using a 3 M KCl reference electrode (Neher, 1992). All experiments were performed at 36°C. Synaptic transmission was blocked with 5 μM NBQX to block AMPARs, 2.5 μM (R)-CPP to block NMDARs, 5 μM SR94431 to block GABA_A_ receptors, and 1 μM strychnine to block glycine receptors.

For each experiment in **Figure 3-supplement 1**, stable responses to a 200 pA current injection were first obtained in control conditions (1.5 mM Ca^2+^, 1.0 mM Mg^2+^). After 5 minutes, external solution was switched to begin washin. Control and washin solutions always included blockers of synaptic transmission. The following external solutions were used: “Low Ca” (0.1 mM Ca^2+^, 4.0 mM Mg^2+^), “0 Ca” (0 mM Ca^2+^, 4.0 mM Mg^2+^, 1 mM EGTA), apamin (300 nM apamin added to control solution), XE991 (10 μM XE991 added to control solution). For SLICK/SLACK DKO, knockout animals in which Na^+^-activated K^+^ channels and control external solution were used. Washin solutions were allowed to equilibrate for at least 15 minutes before recording responses to current injections.

### Analysis

Recordings were obtained using Multiclamp 700B (Molecular Devices), sampled at 50 kHz and filtered at 4 kHz and collected in Igor Pro (Wavemetrics). Data were analyzed using custom-written scripts in Matlab (Mathworks). Instantaneous firing rates were calculated as the reciprocal of the interspike interval. In Igor Pro, decay times for firing rate were determined from an exponential fit to the instantaneous firing frequency during depolarizations. All summary data are shown as means ± SEM unless otherwise indicated. Before comparing adaptation ratio, half-decay time, peak firing, steady-state firing, AHP amplitude, and AHP half-decay time between control and experimental conditions in **Figure 3 – figure supplement 2**, a Shapiro-Wilk test with significance level of 0.05 was used to test whether data were normally distributed. Most data were found to be normally distributed and subsequently compared to their baseline values obtained before the washin by a two-tailed unpaired Student’s *t*-test. In all cases, the threshold for statistical significance was set at p < 0.05. Control (baseline) steady-state firing rates in **Figure 3 – supplement 2** were found to be asymmetrically distributed and subsequently compared to steady-state firing rates of the experimental conditions using non-parametric Wilcoxon signed rank test.

---

**Figure 3 – figure supplement 1.**
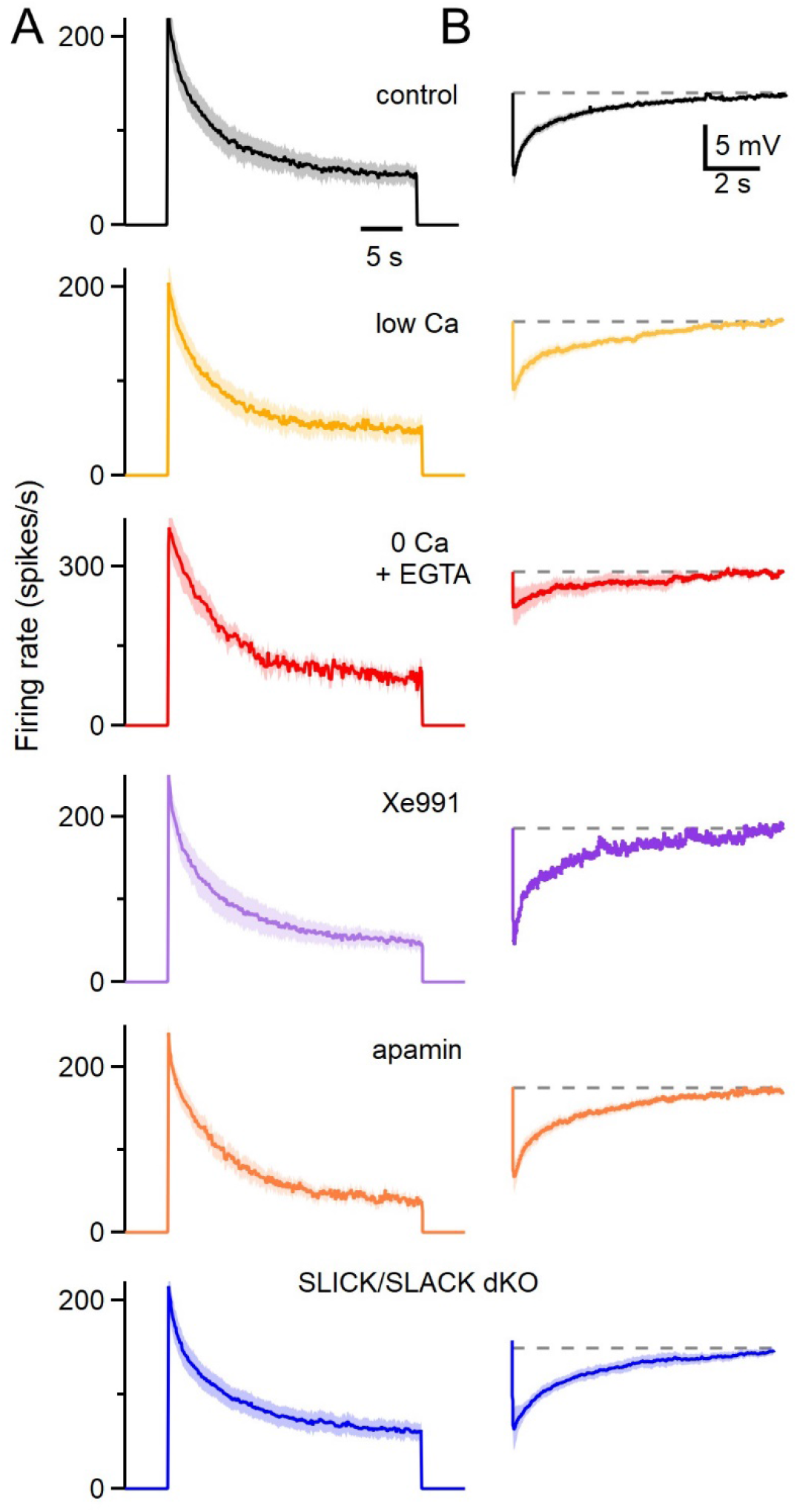
The calcium-dependence and pharmacological sensitivity of spike-frequency adaptation and the subsequent afterhyperpolarization. A. Average instantaneous firing frequencies evoked by 200 pA depolarizing currents for 30 seconds in the indicated conditions. Low Ca solution consisted of 0 mM external Ca^2+^ and 4 mM external Mg^2+^, 0 Ca^2+^ solution consisted of 0 Ca^2+^ and 1 mM EGTA, XE-991 (10 μM, an antagonist of Kv7 (KCNQ) channels), apamin (300 nM, a blocker of SK Ca^2+^-activated K channels), and SLICK/SLACK double knockout animals in which Na^+^-activated K channels have been eliminated. Shading is standard error. B. The average AHPs following the current steps in A are shown for the indicated conditions.

**Figure 3 – figure supplement 2.**
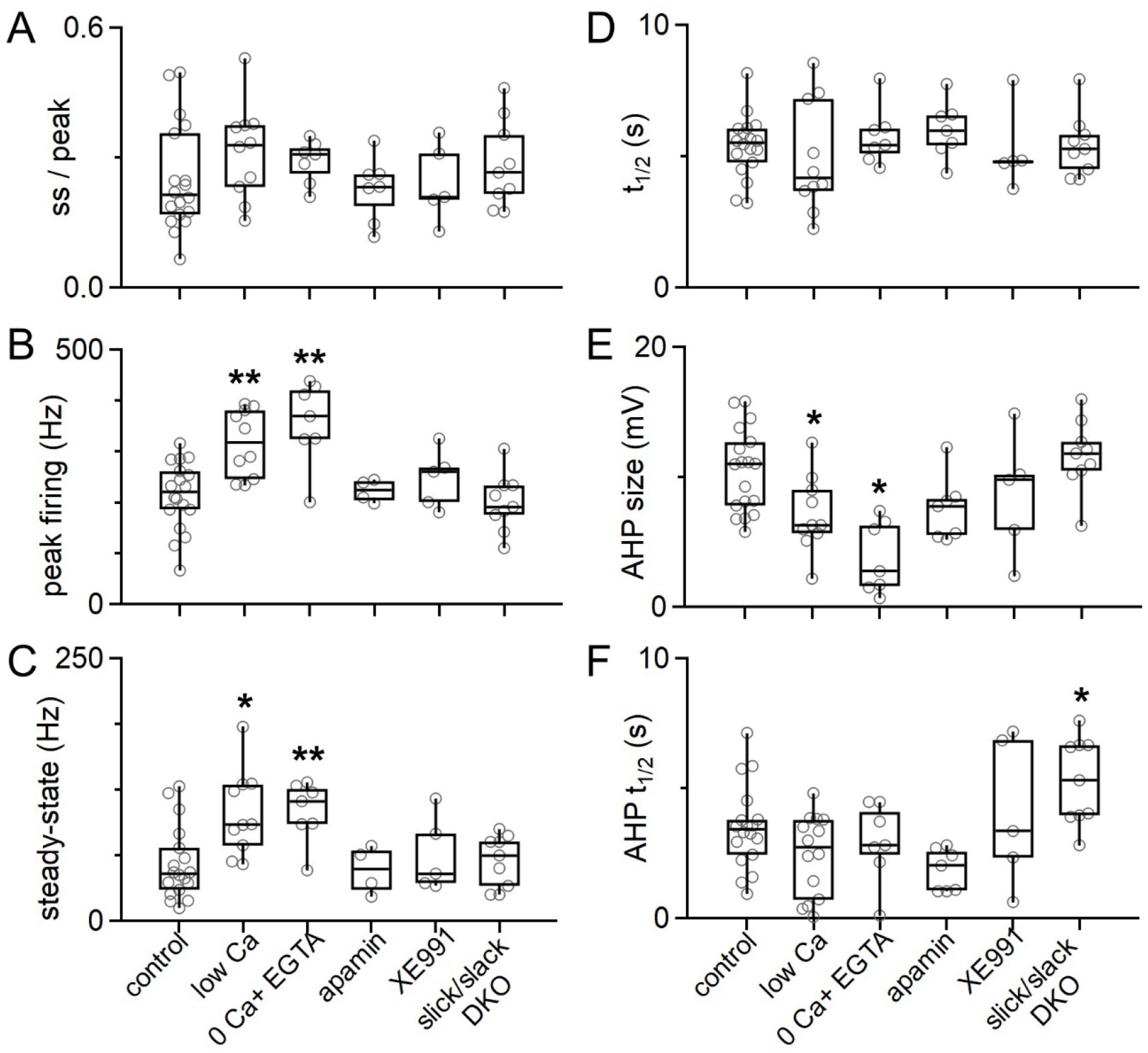
The calcium-dependence and pharmacological sensitivity of spike-frequency adaptation and the subsequent afterhyperpolarization for single cells. A. The steady state / peak firing rates. Conditions are same as indicated in caption for **Figure 3 – figure supplement 1**. B. Peak firing rates occurring during current step. C. Steady state firing rates. D. Decay times for the instantaneous firing frequencies. E. AHP amplitudes. F. The half decay times for AHPs.

## Notes

### Competing Interest Statement

The authors have declared no competing interest.

